# Behavioural and neurogenetic evidence for emotion primitives in the fruit fly *Drosophila*: insights from the Open Field Test

**DOI:** 10.1101/2025.02.28.640745

**Authors:** Yi Lueningschroer-Wang, Emilia Derksen, Hannah Wilde, Maria Steigmeier, Christian Wegener

## Abstract

Emotions, defined as transient states preparing organisms for adaptive responses, are thought to be based on evolutionarily conserved building blocks (“emotion primitives”) that are present across mammalian and other vertebrate species. Whether and to which extend these building blocks are found in insects is largely undefined. In this study, we employ the open field test (OFT) in fruit flies and focus on wall-following and total walking activity as behavioural indicators of emotion-like states.

Wildtype and transgenic flies were subjected to various conditions, including social isolation, starvation, and exposure to anxiolytic substances prior to behavioural testing in the open field test. The results indicate that wall-following and total walking activity are independent measures which are modulated by these conditions, with generally increased wall-following observed under aversive stimuli and decreased values under positive conditions. Notably, the behaviour was consistent across different times of the day, independent of circadian rhythms and largely reversible.

The effects of genetic manipulation of neuromodulatory systems, such as serotonin, dopamine, and neuropeptide F, further supports the role of these pathways in modulating emotion-like states in the fruit fly. Activation of reward-related neurons decreased wall-following, while inhibition increased it, aligning with known effects in mammalian models. Additionally, pharmacological treatments with ethanol and diazepam produced predictable changes in wall-following and total walking activity, reinforcing the validity of the open field test as a measure of emotion-like states in the fly.

The findings suggest that *Drosophila* shares core emotion-like building blocks with vertebrates. Their behaviour in the OFT can serve as a reliable indicator of emotional valence and arousal. This positions *Drosophila* as a powerful genetic model to dissect the neuronal and neurochemical substrates of emotion primitives, shedding light on the evolution of basic emotional processing mechanisms.

## Introduction

Emotions can be defined as transient states that, upon specific events or stimuli, prepare the human organism for adaptive behavioural and physiological responses (Anderson and Adolphs 2014). These states generally involve some form of subjective experience (“feelings”) (James 1884). One example is fear, which triggers specific neuronal and autonomic responses that prime the organism for a range of defensive behaviours, such as freezing and fight or flight, depending on the intensity of the threat stimulus and the likelihood of harm (Roelofs 2017). Threat situations and coupled stimulus-dependent defensive behaviour are encountered by all animals, including humans. It is therefore plausible to assume that basal neuronal circuits and neurochemical brain states underlying fear and other basic emotions in humans are evolutionarily conserved in animals (LeDoux 2012; Anderson and Adolphs 2014), even though it is questionable and difficult to test whether non-human animals experience emotions and conscious feelings. Thus, contemporary concepts of emotions and increasing evidence suggest that basal building blocks in form of neuronal and bodily states underlying emotions in humans (“emotional processing” (LeDoux 2012), “emotion primitives” (Anderson and Adolphs 2014)) can also be found in animals (Damasio 1999; de Waal 2011; LeDoux 2012; Anderson and Adolphs 2014; Zych and Gogolla 2021). These concepts allow the use of animal models, including genetic model organisms such as the mouse, to dissect the neuronal underpinnings of these basal circuits and brain-body states, which are still poorly defined.

Potentially, *Drosophila* may serve as a valuable model for investigating the neuronal and physiological mechanisms of “emotion primitives” (Anderson and Adolphs 2014; Perry and Baciadonna 2017; Gu et al. 2019) due to its sophisticated genetic tools and available whole-brain connectomes (Scheffer et al. 2020; Schlegel et al. 2023). In fact, a range of fly studies have already been conducted to elucidate neuronal mechanisms and neuromodulators underlying specific emotion-related behaviours like aggression (e.g. (Hoyer et al. 2008; Zhou et al. 2008; Williams et al. 2014; Asahina et al. 2014; Jung et al. 2020), or fear-like defensive behaviour (e.g. (Gibson et al. 2015a; Mohammad et al. 2016; Zacarias et al. 2018)). Moreover, evidence of negative cognitive bias, a characteristic of specific affective states, has been reported for another insect, the honey bee (Bateson et al. 2011).

A widely used test for negative (defensive) emotion-like states in rodents is the open field test (OFT). Typically, animals are placed inside an open arena, and the ratio of the time spent in the central vs. peripheral zone is measured and used to assess fear-like behaviour. Increased aversion of the center – commonly referred to as thigmotaxisis usually interpreted as a sign of an increased anxiety/fear-like defensive state (Walsh and Cummins 1976; Simon et al. 1994; Belzung and Griebel 2001; Crusio 2001). Similarly, humans with elevated anxiety sensitivity or patients with anxiety disorders show increased time spent along the green wall of a natural open field (Walz et al. 2016). Outside of a fear-related context, the OFT is used to study exploratory behaviour (e.g. (Easton et al. 2003)) and pain-related thigmotaxis (Zhang et al. 2023).

Similar to rodents, fruit flies show increased center avoidance and wall-following (WAFO) in larger open field arenas (Götz and Biesinger 1985; Besson and Martin 2005; Liu et al. 2007; Valente et al. 2007; Soibam et al. 2012), which at least in part represents exploration of the boundary rather than thigmotaxis (Soibam et al. 2012). A pioneering study from the Claridge-Chang laboratory employed a small open field arena to provide evidence for ancient fear-related pathways that are conserved between mice and flies (Mohammad et al. 2016). Specifically, they found that manipulation of serotonin signaling, anxiolytic treatment with diazepam as well as stress produced remarkably similar effects on WAFO in the OFT as in mice. Thus, *Drosophila* wall following resembled the characteristics of anxiety-related behaviour in rodents, suggesting that WAFO in open field behaviour can be used as a behavioural proxy of a fear-like state in flies.

These findings made us wonder whether WAFO and locomotor activity in the OFT can be used as more general behavioural proxies for emotion-like states beyond fear in *Drosophila*. To investigate this, we subjected different wildtype and transgenic flies to a number of different negative and positive conditions and treatments prior to automated behavioural tracking in the OFT. We conceptually advanced the analysis of OFT behaviour over earlier studies in flies by integrating total walking activity (TOWA, a potential measure of arousal) as a second and independent dimension equal to WAFO.

Across a range of aversive and appetitive conditions, we found that flies show systematic shifts in WAFO and TOWA in the OFT. These effects are reversible, dependent on the genetic background but largely independent of circadian activity and can be altered by modulatory neurons. Collectively, our results show that measuring WAFO and TOWA in a small open field arena provides a useful, low-dimensional behavioural readout of different core emotion-like states in flies beyond the previously reported fear-related context. Hence, this *Drosophila* framework can offer a complementary way to study emotion primitives that allows to employ the available genetic and connectomic tools of the fly to study the neuronal and neurochemical substrates of “emotion primitives”.

## Results

### WAFO and TOWA reversibly correlate with aversive signals in Drosophila

When flies are placed into a larger open field arena, they show a high initial locomotor activity which is strongly dependent on the diameter of the arena and declines quickly over the first minutes to reach a steady activity level (Connolly 1967; Liu et al. 2007). Importantly, Liu et al. (2007) found that this exploratory driven initial activity is completely lost in a smaller circular arena of 6.6 cm diameter, while it was measurable over 3 minutes in arenas of 17.6 and 28.5 cm size. We therefore reasoned that an arena with a diameter smaller than 6.6 cm is best suited to test for the effect of exogenous stimuli on the internal state of the fly.

We first compared activity levels across three smaller arena sizes (5.8 cm, 2.2 cm, and 1 cm in diameter) and chose a circular design to further reduce assay complexity (Martin 2003) compared to a square design as used by Mohammad et al. (2016). As expected, we did not observe a significant initial activity peak within the first minute relative to later time points in all arenas (Figure 1 – figure supplement 1A), although the activity during the first minute was slightly but insignificantly increased in the 1 cm arena. Moreover, the activity level was stable over the 10 min of recording (Figure 1-figure supplement 1B-B’’) and TOWA was similar independent of the arena diameter (Figure 1-figure supplement 1C). We further found that a mechanical stimulus (shake) induced increased wall following behaviour (WAFO, quantified by distance to the center) in all three arena sizes (Figure 1-figure supplement 1D-D’’). TOWA was also increased so the effect was not significant for the 2.2 cm arena (Figure 1-figure supplement 1D-D’’).

**Figure 1:**
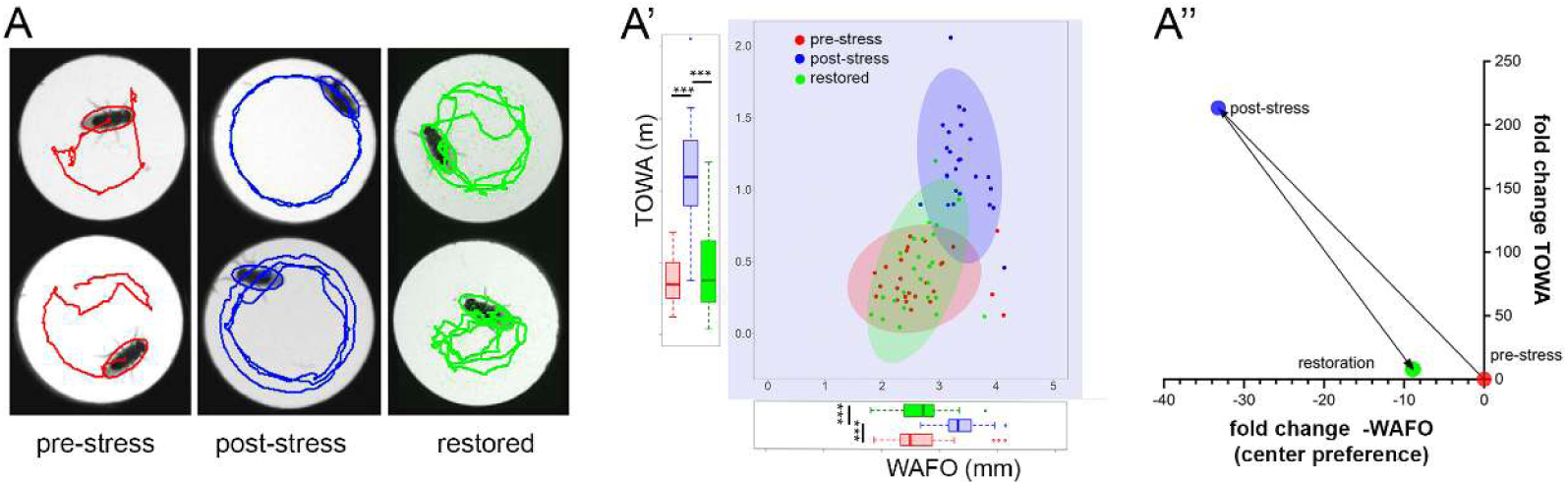
Wall-following behaviour (WAFO) and total walking activity (TOWA) in the OFT upon exposure to multiple stressors. **A)** Typical examples of walking trajectories in the glass-covered arena before experiencing multiple stressors (red), directly after exposure to multiple stressors (blue, one day of social isolation, starvation and sleep deprivation) and after restoration (green, 1 day on food together with other flies of mixed sex and without shaking) as automatically tracked by Ctrax. **A’)** WAFO and TOWA values increase significantly after experiencing multiple stressors and decrease back to pre-stress levels after restoration. **A’’)** The median values of the data in A) were normalised to the median pre-stress value (red dot) and plotted as fold change compared to the pre-stress levels as the fold change of - WAFO on the X-axis, and fold change in TOWA on the Y-axis.

Due to the similarities in the OFT behaviour in the smaller arenas, we decided to use an array of circular arenas with a small diameter (1 cm) which allowed to track 24 flies simultaneously with one camera as well as minimised the initial exploratory activity component (Meehan and Wilson 1987). In addition, a 1 cm square arena had successfully been used to study “fear-like” defensive behaviour in the fly (Mohammad et al. 2016). For our experiments, we placed a single male fly into each arena and automatically tracked its behaviour for 10 minutes using Ctrax (Branson et al. 2009). The 10 min time window was selected based on one-hour recordings, which showed that WAFO as well as TOWA remained relatively stable across 10-minute intervals (Figure 1-figure supplement 1E-E’). In contrast to this naïve response, the stress-induced response in WAFO and TOWA appears to be transient ((Figure 1 – figure supplement 2).

In a first set of experiments, we tested 7 days old unmanipulated and unconditioned male CS flies in the arena, which walked close to the wall, but also crossed the center (Figure 1A). We then removed the flies from the arena and kept them socially isolated under starvation on pure agarose on a vibrating shaker (420 rpm) for one day. Afterwards, we observed a significantly increased WAFO and TOWA when retested in the OFT (Figure 1A-A’F). The males were then grouped together in a fresh vial on standard medium without vibration, and twenty virgin females were added. After one day of restoration, the male flies were retested in the OFT and showed restored pre-stress WAFO and TOWA values (Figure 1 A-A’).

To better visualise the behavioural changes, we normalised the individual values for WAFO and TOWA to the median pre-stress values. We then plotted the resulting relative median values for the whole group, with the X-Y axis representing the respective fold change of WAFO/TOWA compared to the pre-stressed condition (Figure 1A’’, the sign of the fold change of WAFO was reversed to reflect that center preference decreases with increasing strength of aversive signals). Due to normalisation, the median pre-stress value now centers at the axis origin, and the fold-change of -WAFO and TOWA can obtain a positive or negative value (Figure 1A’’). After experiencing stress, the values moved in the direction of an increased WAFO and increased TOWA (Figure 1C). Control experiments, in which CS males were tested over three days without exposure to stressors, showed no significant changes in either WAFO or TOWA (Figure 1-Figure supplement 1F-F’), indicating the absence of a habituation effect.

To evaluate the effect of single stressors, seven days old male naïve flies were tested pre-treatment on day 1, and then either kept i) isolated (Figure 2A-A’), ii) starved (Figure 2B-B’), iii) sleep-deprived (Figure 2C-C’), iv) sex-deprived as virgins by female rejection (Figure 2D-D’), or v) sex-deprived after mating in a group without females (Figure 2E-E’) over two to three days (day 2-4). Afterwards, they were re-tested for their behaviour in the OFT. At the end, flies were allowed to restore from stress by enabling social contact, feeding, mating and sleep according to the previously exerted stress for one day (day 4 or 5), followed by a final test in the OFT. As before, the unconditioned pre-stressed flies defined the baseline value to normalise the results. After social isolation for one and two days (Figure 2A-A’), or two days of starvation (Figure 2B-B’), flies showed significantly increased TOWA. WAFO increased as well after two days of social isolation (Figure 2A-A’), and after one and two days of starvation (Figure 2B-B’). These changes in behaviour further increased with four days of social isolation, or two days of starvation (Figure 1A-B’). The strong increase in TOWA from day 1 to day 2 under starvation can at least partially be explained by increased AKH/octopamine-dependent starvation-induced hyperactivity (Lee and Park 2004; Isabel et al. 2005; Yang et al. 2015; Yu et al. 2016). Upon restoring social contact and providing food, TOWA and WAFO shifted back towards the pre-stress levels which were, however, not fully reached at least for WAFO (Figure 2A-B’). Similarly, when either sleep-deprived during the day including siesta time (Figure 2C-C’), or sex-deprived as virgins due to being rejected by mated females for one day (Figure 2D-D’), flies increased both TOWA and WAFO. These changes became again more pronounced over time with continued sex deprivation like the changes seen after social isolation and starvation (Figure 2D-D’). In line, extending sleep deprivation into the night resulted in an increase of WAFO, while TOWA decreased possibly due to exhaustion from night sleep deprivation. After restoration, WAFO for both sleep- and sex-deprived flies shifted towards pre-stress levels, while TOWA remained significantly higher compared to the pre-treatment level (Figure 2C-D’). When mated adult flies were sex-deprived by keeping them in male-only groups, TOWA and WAFO levels (Figure 2E-E’) changed qualitatively like what was observed for sex-deprived virgin males that were rejected by mated females (Figure 2D-D’). Notably, restoration by mating with a virgin female decreased both WAFO and TOWA (Figure 2E-E’). One difference in TOWA levels between the restored sex-deprived males is that the mating-rejected virgin males (Figure 2D-D’) were sexually naive, while the mated males kept separated from females (Figure 2E-E’) already had the opportunity to mate prior to the test. The increased TOWA levels of the virgin males after first mating during restoration may thus indicate an increased motivation to look for new mating partners due to a stronger activation of a neuronal reward circuit compared to mating-experienced males. Combined, the results for the different single stressors show that stress in general leads to increased TOWA and WAFO levels, while restoration from stress in general reverts both values to pre-stress levels.

**Figure 2:**
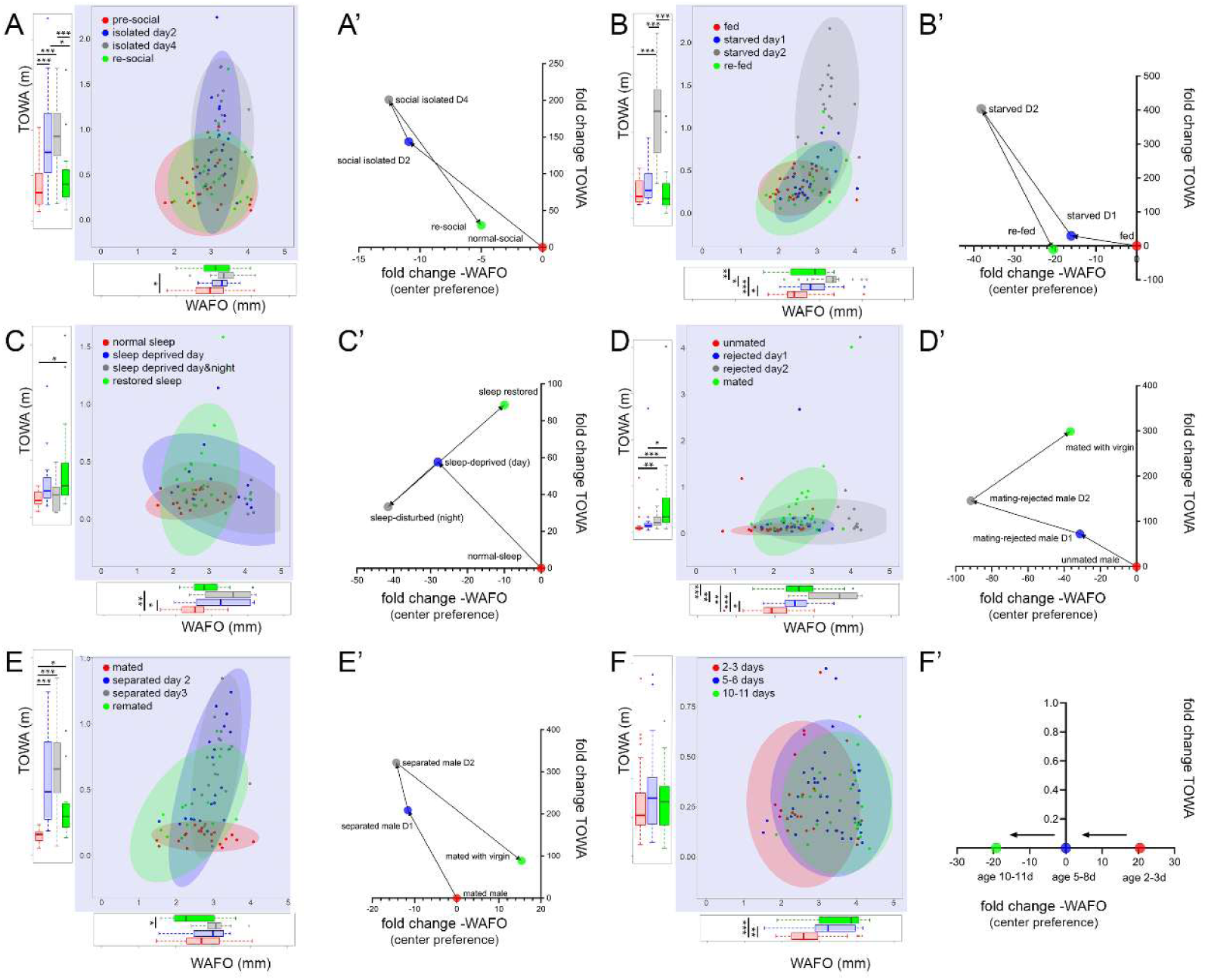
Wall-following behaviour (WAFO) and total walking activity (TOWA) with different stressors. Raw data are shown in X), while X’) shows the median after normalisation to the pre-treatment condition, plotted as fold changes. **A-A’)** Effect of social isolation. **A)** Flies were tested in the OFT, then socially isolated and re-tested on day two and four, and then regrouped and tested again. Both WAFO and TOWA are strongly affected by social isolation. **A’)** With increasing social isolation, flies walk increasingly more and closer to the wall. After restoration of social contact, data points shifted back in direction to pre-stress levels. **B-B’)** Effect of starvation. **B)** Flies were tested in the OFT, then starved and tested after one and two days in starvation and then retested again after one day on food. Both WAFO and TOWA are strongly affected by starvation. **B’)** Starvation for one and two days increasingly leads to higher TOWA and WAFO levels. After re-feeding, levels shift back towards pre-starvation levels. **C-C’)** Effect of sleep deprivation. **C)** Flies were tested in the OFT, then mechanically sleep deprived and tested in the middle of the light phase (ZT7) and at the beginning of the dark phase (ZT13) of day 1 during sleep interruption. Following a day of mechanically undisturbed sleep, flies were then retested. WAFO but not TOWA was strongly affected by sleep deprivation. **C’)** Sleep deprivation for the first half or the whole photophase lead to increased WAFO but had negligible effect on TOWA. After restoration of sleep, WAFO levels approached pre-stress levels with strongly increased locomotor activity. **D-D’)** Effect of mating rejection on unmated males. **D)** Unmated male flies were tested in the OFT and then paired with mated females. After being mating-rejected from these females for an hour each on day 1-2, male flies increasingly shifted towards higher TOWA and WAFO levels. After restoration by a successful mating on day 3, a further increase in TOWA but a partial decrease in WAFO was observed. **D’)** After being rejected by females on day 1 and 2, males become increasingly active and show increased WAFO compared to unmated males. After restoration of courtship, the males showed a further increase in locomotor activity and a decrease in WAFO levels towards the pre-stress condition. **E-E’)** Effect of sexual deprivation on mated males. **E)** Male flies were mated at day 1 and afterwards tested in the OFT, then sexually deprived and tested on day 2 and 3 of sexual deprivation. **E’)** This treatment led to a strong and increasing shift towards higher TOWA levels, while WAFO did not change significantly. After restoration by a successful mating on day 3, TOWA levels dropped back and WAFO increased compared to 2 days of sexual deprivation. **F-F’)** Effect of age. **F)** Different groups of male flies were tested in the OFT at an age of either 2-3, 5-6 and 10-11 days. **F’)**This treatment led to a strong and increasing shift towards lower WAFO levels, while TOWA did not change significantly.

To test for the effect of age, we analysed the open field behaviour of unconditioned young flies 2-3 days, 5-6 days, and 10-11 days after eclosion (under our conditions, flies typically reach an age of 50+ days). We found no effect in TOWA between these age groups (Figure 2F-F’), indicating that the flies are physiologically of a similar (young) age. In contrast, we observed a significant increase of WAFO in the older groups compared to 2-3 days old flies (Figure 2F-F’), which suggests that older flies are more defensive than young flies. Significant differences in WAFO between 5-6- and 10-11-day old flies were not observed, which validates our standard use of 5–11-day old flies for the OFT. These age-related changes in WAFO suggest that either age-dependent experience or (brain) maturation during early life alter open field behaviour.

### Aversive stimuli and anxiolytic substances influence the open field behaviour of flies similar to mice

We next ask whether aversive stimuli connected to pain in rodents lead to similar behavioural changes in flies. For that, we applied i) a mild mechanical shock (5x tapping) to induce a startle response, ii) a heat pulse of 37°C during the OFT, and iii) ten brief consecutive electric foot shocks. Flies were tested for their open field behaviour before and during or after the treatment. A mild mechanical shock significantly increased WAFO compared to pre-stimulus conditions (Figure 3A-A’). This effect was intensified after the mechanical shock was repeated (Figure 3A-A’), along with a significant increase in TOWA. Similarly, a 37°C heat shock during the OFT increased both WAFO and TOWA (Figure 3B-B’). Moreover, a medium aversive electric foot shock (EFS) resulted in increased WAFO and TOWA (Figure 3C-C’). We next repeated the experiment with EFS with a new set of flies (Figure 3D-D’). As before, TOWA was increased, while WAFO reached from medium to elevated levels after EFS (Figure 3D-D’). The day after, we first retested the flies which showed that WAFO and TOWA were fully restored to pre-shock conditions. Afterwards, we applied a second EFS. As expected, this resulted in increased WAFO value similar to the EFS on day 1. Strikingly, however, the increase in TOWA was lost after the second EFS (Figure 3D-D’), reminiscent of the learned “helplessness” state of flies after EFS (Batsching et al. 2016).

**Figure 3.**
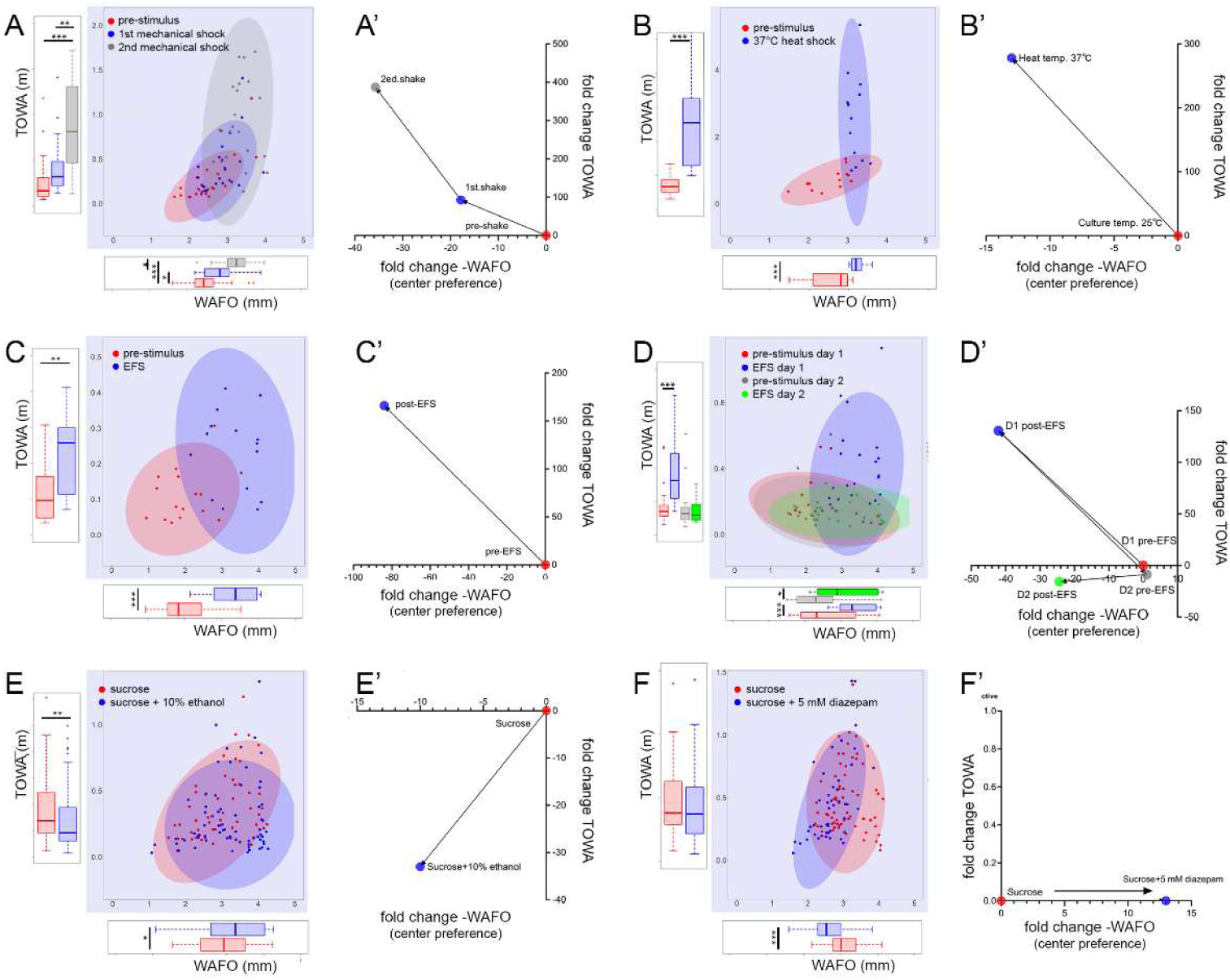
Aversive stimulation and pharmaceutical treatment consistently alter open field behaviour. Raw data are shown in X), while X’) shows the median after normalisation to the pre-treatment condition, plotted as fold changes. **A-A’)** Effect of mechanical shocks (tapping). **B-B’)** Effect of a 10 min heat-shock of 37°C during the OFT and **C-C’)** an electric foot shock (EFS) prior to the OFT increased TOWA and WAFO compared to pre-stimulus conditions. **D-D’)** The effect of an EFS given on two consecutive days confirmed the increase of WAFO and TOWA after the first shock, while the second shock had a reduced effect on WAFO and no effect on TOWA. **E-E’)** Effect of 10% ethanol. Feeding on 10% ethanol in sucrose induced a mild increase in WAFO compared to sucrose alone, while TOWA was weakly reduced. **F-F’)** Effect of 5 mM diazepam. Feeding on 5 mM diazepam plus sucrose induced a stronger increase in WAFO compared to sucrose alone, while TOWA was unaffected.

The OFT is often used to screen for effects of drugs, including ethanol and diazepam, on pain-induced avoidance and fear-like behaviour in rodent models (Prut and Belzung 2003; Knight et al. 2020; Zhang et al. 2023). To assess whether flies react similarly, we tested the effect of ethanol (which has different dose-dependent effects on locomotor activity in flies (Singh and Heberlein 2000)) and diazepam (a benzodiazepine and prototype anxiolytic drug). Using a capillary feeder (CAFE) assay to quantify food intake (Ja et al. 2007), we first supplied flies for 24 hours with 10% ethanol mixed with sucrose and then tested flies that had consumed at least 1µl. After ethanol consumption, flies showed weakly increased WAFO, and a weakly decreased TOWA compared to controls that consumed sucrose only (Figure 3E-E’). Thus, feeding on 10% ethanol had a weak sedative but no “anxiolytic” effect on flies under our conditions. Next, we kept flies in vials on 2% coloured agarose mixed with either 4% sucrose (control) or 4% sucrose plus 5 mM diazepam. Food consumption was checked by the appearance of colour in the gut of the flies. Flies that had consumed sucrose plus diazepam-containing food showed a significantly decreased WAFO while TOWA was unaffected compared to controls that had consumed only sucrose (Figure 3F-F’). This indicates that, unlike 10% ethanol, diazepam has an “anxiolytic-like” effect and confirms that the “anxiolytic” effect of a diazepam-ethanol solution described in an earlier study (Mohammad et al. 2016) was indeed due to the benzodiazepine.

### Genetic manipulation of neuromodulator signalling affects WAFO rather than TOWA

The results above show that stress and aversive signals trigger a similar open field behaviour in flies as reported for rodents. In mammals including humans, mood, anxiety and fear are under neuromodulatory control by biogenic amines and neuropeptides (Rana et al. 2022; Pourhamzeh et al. 2022). In particular, serotonin (5-hydroxytryptamine, 5-HT), dopamine as well as neuropeptide Y (NPY) signalling are known to strongly affect the open field behaviour of rodents ((Simon et al. 1994; Broderick and Phelix 1997; Prut and Belzung 2003). We therefore asked whether manipulating homolog neurochemical systems in the fly brain also yields similar effects in the open field behaviour as found in mice. First, we revisited the role of serotonin signalling by repeating previous experiments by Mohammad and colleagues (2016). We conditionally down-regulated (by RNA interference) or overexpressed the *Drosophila* serotonin transporter (dSerT) using a temperature-inducible (29°C) pan-neuronal *nSyb tubGal80^ts^* Gal4 driver line based on the TARGET system (McGuire et al. 2003). Control and experimental flies were pre-incubated at either 18°C or 29°C prior to the OFT assays conducted at 20°C. The pre-incubation temperature itself had no effect on the respective GAL4 driver and UAS responder controls (Figure 4A-D). Overexpression of dSerT at 29°C significantly reduced WAFO (Figure 4A) in line with the results of Mohammad et al. (2016). RNAi-mediated down-regulation of SERT at 29°C, however, also resulted in reduced WAFO (Figure 4A), contrary to the results by Mohammad et al. (2016). TOWA was not significantly affected by either temperature-dependent conditional downregulation or overexpression of dSerT (Figure 4-supplementary Figure 1A). Directly after the OFT, the flies received an electrical foot shock (EFS) in the arena and then were tested a second time. This treatment significantly increased both WAFO (Figure 4-supplementary Figure 1C) and TOWA (Figure 4-supplementary Figure 1D) in controls and flies with down-regulated SerT expression. Yet, flies with *SerT* overexpression showed a significant increase only for TOWA (Figure 4-supplementary Figure 1D). Moreover, decreased or increased *dSerT* expression at the permissive temperature led to a significantly reduced post-EFS increase in WAFO compared to the non-permissive temperature (Figure 4B). Next, we used the *GMR58E02-Gal4* line and expressed a temperature-sensitive form of dynamin encoded by *shibire* (Shi^ts^) (Kitamoto 2001) to specifically and conditionally suppress neuronal transmission of the dopaminergic protocerebral anterior medial (PAM) neurons that project prominently to the medial lobe of the mushroom body (Liu et al. 2012). While the PAM neurons are an inhomogeneous group of neurons in adult flies, their *en masse* activation (Liu et al. 2012; Burke et al. 2012) or disinhibition (Yamagata et al. 2021) signals reward, and their thermogenetic or optogenetic activation induces valence-driven behaviour such as attraction or aversion (Aso et al. 2014; Mohammad et al. 2024). Specific silencing of neuronal transmission at 29°C in PAM neurons significantly increased WAFO compared to the same genotype tested at 18°C (Figure 4C), while temperature itself had no effect in controls (Figure 4C). As in the dSerT experiment, we found no change in TOWA after suppression of PAM neuron signalling (Figure 4-supplementary Figure 1B). Next, we activated the PAM neurons thermogenetically at 29°C via the temperature-sensitive channel TRPA1 prior to the OFT. This resulted in a significant decrease in WAFO (Figure 4C) compared to temperature controls. Interestingly, this was accompanied by a significant decrease in TOWA (Figure 4-supplementary Figure 1B).

**Figure 4.**
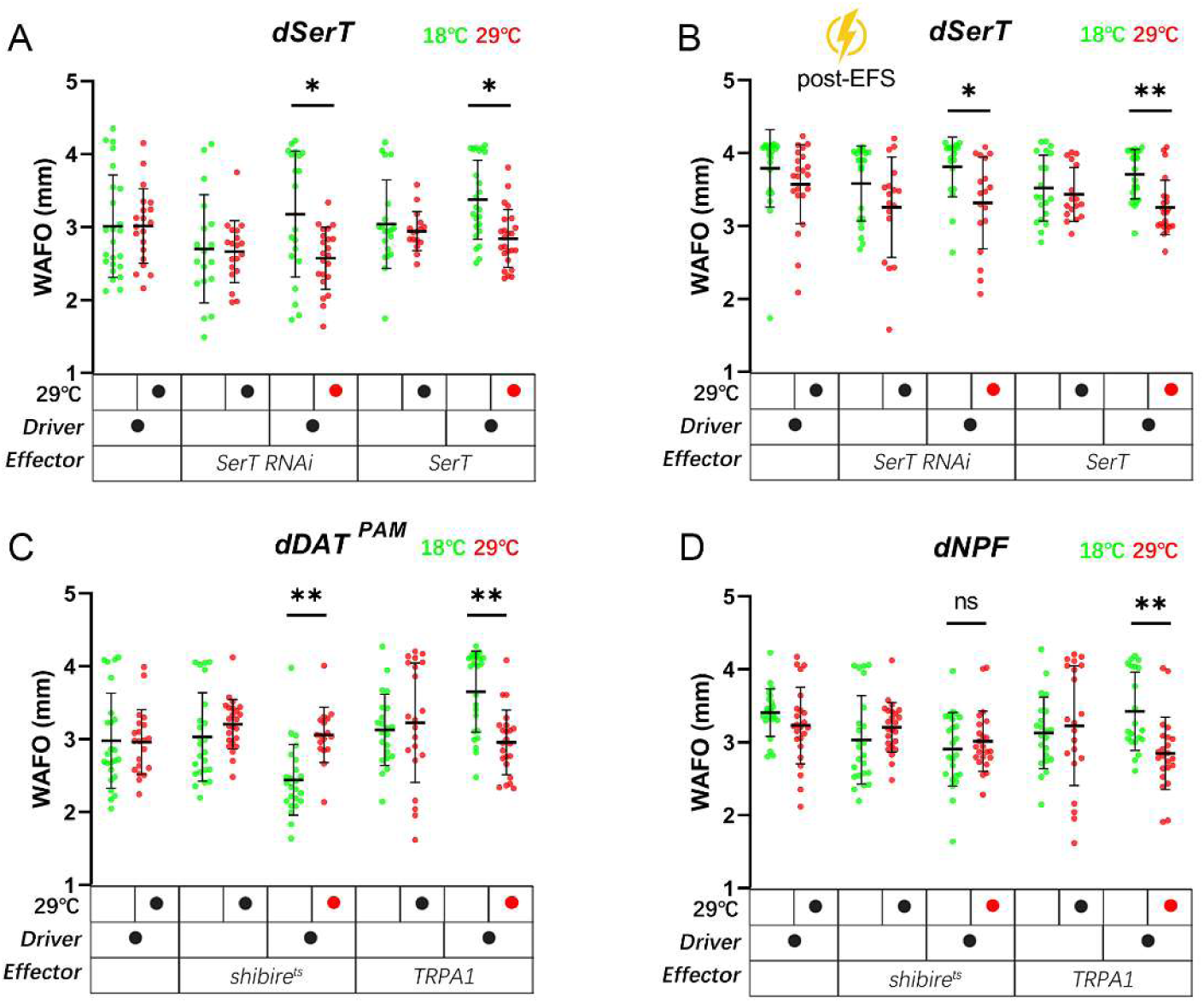
Genetic manipulation of aminergic and peptidergic signalling affects open field behaviour. **A)** Flies with temperature-induced (29 °C, red) conditionally reduced serotonin transporter expression (nSyb tubGal80^ts^ x SerT RNAi) or UAS-mediated SerT overexpression (nSyb tubGal80^ts^ x SerT) showed significantly reduced WAFO compared to flies at the non-inducing condition (18 °C, green), while WAFO was unaffected by temperature in the genetic controls. **B)** Similar experiment as in A) but after an electric foot shock (EFS) showed a reduced but still significant decrease in WAFO after manipulation of SerT expression. **C)** Thermogenetic suppression of dopaminergic reward neurons of the PAM cluster significantly increased WAFO compared to their temperature control, while thermogenetic activation of the same PAM neurons decreased WAFO value compared to their temperature control. Genetic controls were not significantly affected by temperature. **D)** Thermogenetic activation of reward-mediating NPF-positive neurons significantly decreased WAFO compared to temperature controls, while thermogenetic silencing had no effect. Temperature did not significantly affect WAFO in the genetic controls.

Neuropeptides represent the most diverse group of neuromodulators, but -unlike biogenic amines-have so far not been tested for their effect in the OFT in the fly. To broadly assess the effect of neuropeptides on WAFO and TOWA, we tested *silver* (*svr*) mutant flies (Sidyelyeva et al. 2006, 2010) which are impaired in neuropeptide production (Pauls et al. 2019). *Svr* encodes for carboxypeptidase D (CPD) and *svr* mutants where rescued by daily heat shock-driven expression of a CPD transgene (Sidyelyeva et al. 2010). Prior to testing, flies were allowed to deplete their neuropeptide stores for one to three days in the absence of a heat-shock. These acutely neuropeptide-depleted s*vr* mutants showed a significantly decreased WAFO compared to the genetic controls (Figure 4-supplementary Figure 1E), while TOWA remained unaffected (Figure 4-supplementary Figure 1E’). This result suggests that neuropeptides may be involved in regulating emotion primitives. While the net effect of acute neuropeptide depletion obviously is a decreased WAFO, individual neuropeptides may have differential effects, and some may have rather an anxiolytic effect. Mammalian NPY as well as its fly homolog neuropeptide F (NPF) are key peptides in the regulation of feeding and mating motivation, as well as reward (Krashes et al. 2009; Nässel and Wegener 2011; Loh et al. 2015; Comeras et al. 2019; Ryvkin et al. 2024). In mammals, NPY signalling has an inhibitory effect on fear and anxiety in order to facilitate foraging and other rewarding behaviours (van den Burg and Stoop 2019; Comeras et al. 2019). In flies, thermogenetic activation of NPF neurons is perceived as rewarding as shown by olfactory conditioning (Shohat-Ophir et al. 2012). Moreover, mating-rejected isolated males show significantly decreased NPF transcript and peptide levels compared to virgin or mated controls (Shohat-Ophir et al. 2012). Therefore, we first blocked neuronal transmission specifically in the NPF neurons using *shi^ts^*, yet this did neither affect WAFO nor TOWA (Figure 4D, (Figure 4-supplementary Figure 1F). Thermogenetic activation of NPF-positive neurons via TRPA1 led to a significant decrease of WAFO (Figure 4D) but did not alter TOWA (Figure 4-supplementary Figure 1F).

Collectively, these findings show that the open field behaviour of *Drosophila* is influenced by neuromodulators for which the mammalian homologues are known to affect emotional states and open field behaviour.

### In the OFT, wild-caught Drosophila melanogaster strains are more defensive and aroused than laboratory wildtype strains

The results above show that fruit flies in the OFT react similarly to stressors and impaired neuromodulatory signalling as rodents for which the open field behaviour can be linked to fear-like states. Like other emotions, fear and anxiety are products of evolution, thought to promote the adaptation of behavioural and physiological responses to the environmental condition, e.g., the presence of a predator or other dangerous situations. However, all experiments so far were conducted with fly strains that for many generations are kept in the laboratory without predators, with *ad libitum* access to food, and with extremely limited possibility to spatially escape from aggressors. We therefore wondered to which degree the observed OFT responses are representing natural responses, and compared the unconditioned open field behaviour among three laboratory wild type strains (OregonR (OrR), CantonS strain (CS), and CantonS flies that were outcrossed against a *period^01^*strain for 110 generations (CS^110^) (Horn et al. 2019)), as well as wild-caught strains from Hubland-Würzburg/Germany (Deppisch et al. 2022) Munich/Germany and Akaa/Finland (coll: John Parsch and Maaria Kankare within the DrosEU consortium (Kapun et al. 2020)). To cover the natural variation within the wild-caught strains, all experiments were conducted with the F1 outcross of isofemale lines from the respective population. To account and test for possible time-dependent effects, an OFT was performed every six hours during a day in a light-dark cycle of 12:12. Daily rhythmicity or significant day-dependent changes in WAFO were not observed in the tested strains (ANOVA, p > 0.05, Figure 5A-B) except for the CS^110^ strain (ANOVA, p<0.01). We infer from this that the naïve WAFO behaviour in the OFT is independent of the time-of-day and the circadian clock. For TOWA, only the Finnish (ANOVA, p<0.0001) and the CS^110^ strain (ANOVA, p<0.05), but not the other strains showed significant day-dependent differences in TOWA (Figure 5 A-B, Suppl. Table 1). These differences, however, did not correlate with the typical morning (around ZT0) and evening peak (around ZT12) in spontaneous locomotor activity in the DAM assay (Figure 5C) which is under circadian control (Schlichting and Helfrich-Förster 2015).

**Figure 5.**
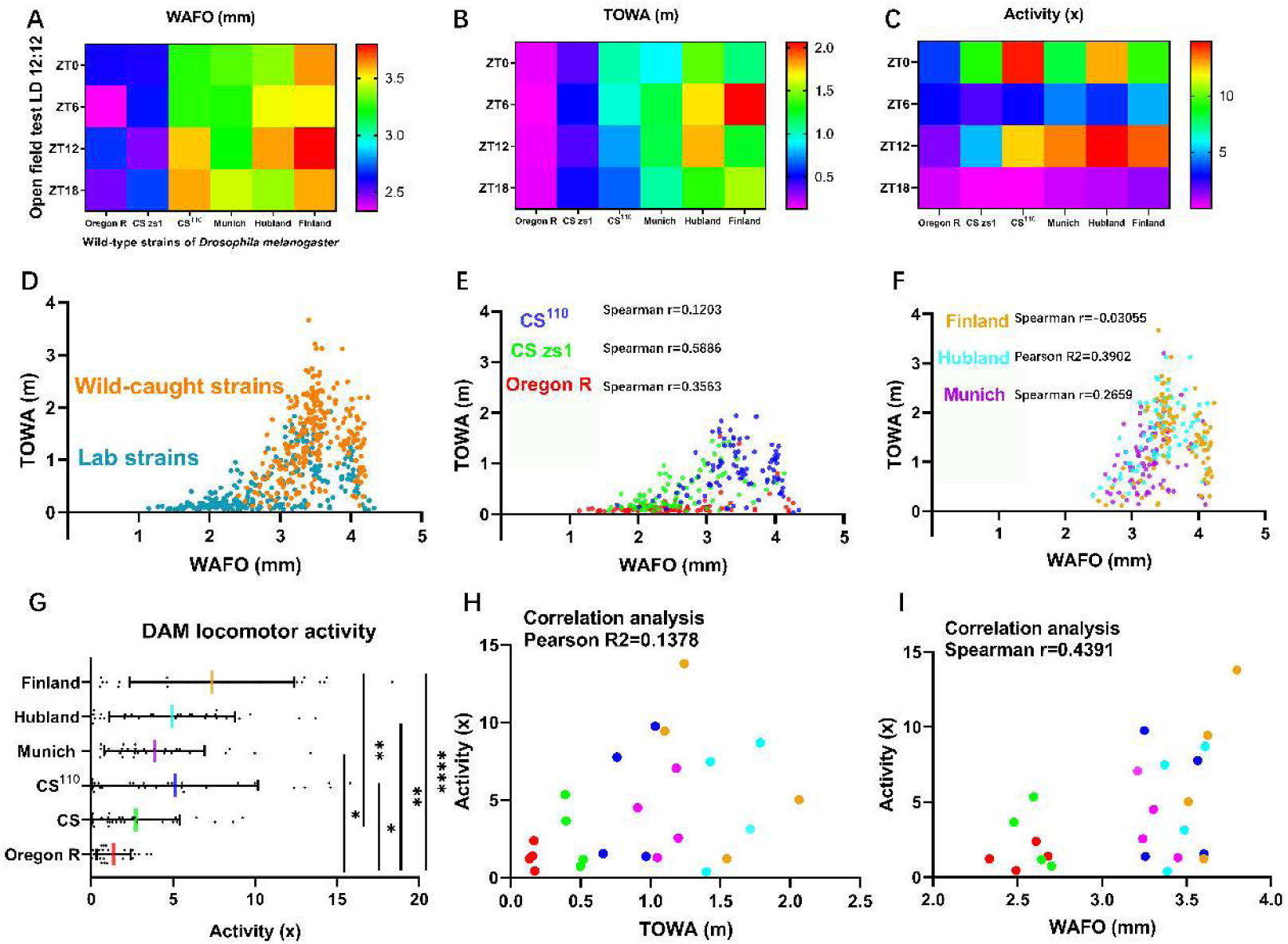
Comparison of WAFO and TOWA and spontaneous activity between naïve laboratory (OrR, CS, CS^110^) and wild-caught Drosophila melanogaster wild-type strains. **A)** Heat map for WAFO values obtained in 6h intervals during the day. WAFO appeared to be much lower in the inbred lab strains (CS, OrR) compared to an outcrossed lab strain (CS^110^) and wild-caught strains. **B)** Heat map for TOWA values obtained in 6h intervals during the day. TOWA appeared to be much lower in the inbred lab strains compared to an outcrossed lab strain (CS^110^) and wild-caught strains. **C)** Heat map of spontaneous locomotor activity as measured in the TriKinetics DAM system, obtained in 6h intervals during the day. Spontaneous locomotor activity appeared lower in the inbred lab strains compared to an outcrossed lab strain (CS^110)^ and wild-caught strains. . All strains show higher activity during the morning (ZT0) and evening (ZT12), in line with a circadian morning and evening peak in locomotor activity. **D)** Scatter plot of the WAFO and TOWA values obtained in the laboratory strains (CS, CS^110^, OrR) and the wild-caught strains. Very low WAFO values were only observed in the laboratory strains. **E)** The inbred laboratory strains CS and OrR but not the outcrossed CS^110^ strain showed a weak linear relationship and positive correlation between TOWA and WAFO. **F)** In the wild-caught strains, WAFO and TOWA were not (Finland, Munich) or only weakly (Hubland) positively correlated. **G)** Spontaneous locomotor activity in the DAM monitors was generally higher in the wild-caught strains than in the laboratory-derived strains. **H)** The spontaneous locomotor activity in the DAM monitors was only weakly correlated with the TOWA in the OFT. **I)** The spontaneous locomotor activity in the DAM monitors was only weakly and non-monotonously correlated with WAFO in the OFT.

Of note, the two inbred lab strains OrR and CS showed significantly lower average WAFO and TOWA values compared to the outcrossed CS^110^ and the wild-caught strains (Kruskal-Wallis with Dunn’s multiple comparison, p< 0.001, Figure 5 A-B, D). Moreover, the average spontaneous activity in the DAM monitor was significantly lower in the inbred CS and OrR strains compared to the outcrossed CS^110^ and the wildtype strains (Kruskal-Wallis with Dunn’s multiple comparison, p< 0.0001, Figure 5C, G). These results could be explained by adaptation of the laboratory strains to a spatially restricted life in small vials with *ad libitum* availability of food, resulting in reduced general locomotor activity and reduced drive to escape.

WAFO and TOWA were at best only weakly correlated (Figure 5 E-F). Similarly, TOWA (Pearson R^2^=0.1378, Figure 5H) and WAFO (Spearman r=0.4391, Figure 5I) correlated only weakly with the spontaneous activity measured in the DAM monitors. Taken together, these findings suggest that WAFO, TOWA and spontaneous activity are largely independent variables.

### Is the OFT response sex-specific and dependent on the mating status?

All experiments so far have been carried out with male flies. To test whether the observed responses are male-specific, we compared the naïve OFT response of mated and unmated CS males and females. Without prior stress treatment, flies showed similar WAFO values independent of sex and mating status (Figure 6A), in line with an earlier study in a larger circular arena (Bath et al. 2020). In contrast, the naïve TOWA response significantly depended both on sex and mating status (Figure 6A’). The low TOWA of mated females compared to virgins was unexpected since mated females are known to increase their spontaneous locomotor activity in a sex-peptide-dependent manner during the photophase (Isaac et al. 2010). This suggests that TOWA in females is largely independent of spontaneous clock-regulated activity, supporting the above findings for males. We next tested the stress response of mated females after receiving a mechanical shock (tapping). Unlike in males (Figure 3A), a single mechanical shock had no effect on WAFO, yet significantly increased TOWA (Figure 6B-B’). A second mechanical shock induced a significant increase in both WAFO and TOWA (Figure 6B-B’). This result suggests that the OFT response is qualitatively similar between the sexes and mating status, while the threshold for stress-elicited responses may differ.

**Figure 6.**
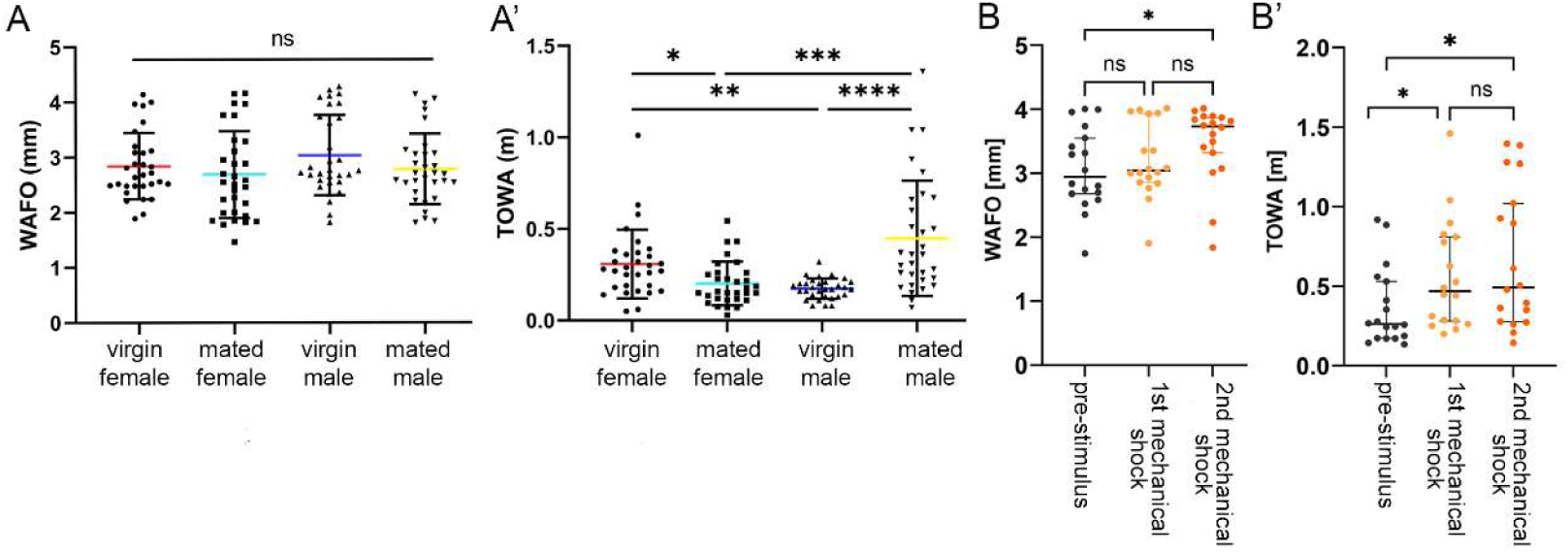
TOWA but not WAFO depends on sex and mating status. **A**) Sex and mating status does not influence wall following in naïve CantonS flies. **A’)** Mating has opposite effects on locomotor activity in the OFT between the sexes. While mated females show significantly reduced TOWA compared to virgins, mated males are significantly more active than unmated males. The active and inactive states between the sexes show similar activity levels. **B)** Mated females show a WAFO response only after repeated mechanical shock (tapping), while TOWA is increased directly after the first mechanical stimulus (B**’**).

## Discussion

The main aim of this study was to evaluate whether wall-following (WAFO) and total walking activity (TOWA) in the OFT can be used to behaviourally assess core dimensions of affect-like states in fruit flies. Our results show that conditions and treatments associated with negative valence in humans or other mammals (including social isolation, starvation, sex or sleep deprivation, heat, mechanical disturbance, electric foot shocks) generally increased WAFO. On the contrary, conditions and treatments associated with positive valence in mammals (including social company, feeding, relieve from stress) generally decreased WAFO. This raises the question whether WAFO can be interpreted as indicating emotional valence in the fly.

### WAFO as an indicator of valence in the fly?

In humans, valence can be defined as an indicator whether something is good or bad/pleasant or unpleasant/helpful or harmful. As such, valence is one key dimension of emotional states, reflecting the organism’s capacity for evaluating stimuli or situations (Barrett 2006). In behavioural terms, positive valence leads to approach behaviour (positive taxis), while negative valence results in withdrawal (negative taxis). In T-maze experiments with *Drosophila* (Tully and Quinn 1985), for example, approach or avoidance of an odour learned to be associated with either reward or punishment is extensively used to dissect the neuronal mechanisms underlying appetitive and aversive memory and valence assignment (e.g. (Keene 2004; Perisse et al. 2013; Aso et al. 2014). One outcome of these and similar studies is that distinct subsets of dopaminergic neurons in the fly brain assign and innately encode positive or negative valence of odors or tastes (Owald and Waddell 2015; Siju et al. 2020). Strikingly, optogenetic activation of a large subset of dopaminergic neurons in the paired-anterior-medial (PAM) cluster (targeted by R58E02-Gal 4 (Liu et al. 2012)) that drives appetitive memory to odors was alone sufficient to elicit attraction in a light-dark optogenetic assay Thermogenetic activation of the R58E02 PAM neuron subcluster also increased attraction to the center in the OFT, while conditional block of their activity increased WAFO. Thus, WAFO-center preference appears to be a suitable behavioural indicator of emotional valence in the fly.

In the study of human emotions, dimensional theories (Russell 1980; Watson et al. 1999; Posner et al. 2005) have gained considerable influence and provide a conceptual framework that is extendable to the study of animals as they rely on few empirically measurable dimensions (Mendl et al. 2010; Anderson and Adolphs 2014). One of the most widely used dimensional theories is the circumplex model that is based on two bipolar dimensions thought to represent two independent neurophysiological systems: arousal (active – inactive) and valence (positive/pleasure-negative/displeasure) (Russell 1980; Posner et al. 2005). Subjective experiences that can be classified by these dimensions are referred to as “core affect”, regarded to be a most basic component underlying emotions (Russell 1980, 2003). Core affect represents the “constant stream of transient alterations in an organism’s neurophysiological state that represents its immediate relationship to the flow of changing events” (Barrett 2006), i.e. changes in reaction to internal and external stimuli. A great advantage of the circumplex model is that the degree of arousal and valence can be obtained by oral report but also by objective physiological and behavioural measurements. This allows to extend the circumplex model to non-human animals where subjective report is absent but behaviour and physiology can still be measured (Mendl et al. 2010). Based on our conclusion that WAFO can be used as a proxy for valence, this raises the question whether TOWA can be used as a proxy for arousal, and hence whether the observed OFT behaviour in the fly can be plotted in a circumplex-like manner.

### TOWA as a proxy for exogenously induced arousal in the fly?

Arousal, the second fundamental axis of the circumplex model in humans, describes the continuous spectrum of physiological and psychological activation, that is, the extent to which an emotional state is experienced as energizing (Russell 1980, 2003; Barrett et al. 2007). Altered locomotor activity is part of an operational definition of arousal (Pfaff and Banavar 2007) and is a hallmark of emotion states across species (Zych and Gogolla 2021). In *Drosophila*, like in other animals, arousal and locomotion are shaped by internal factors such as sleep drive, satiety, emotional state or the circadian clock, as well as by external stimuli including environmental novelty (van Swinderen and Andretic 2003; Van Swinderen and Andretic 2011). For example, elevated locomotor activity in response to a novel or threatening visual stimulus is commonly interpreted as increased arousal and may reflect defensive, emotion-like processes (Gibson et al. 2015b). In our study, we used a small OFT arena to minimise exploratory behaviour (Liu et al. 2007; Soibam et al. 2012). Under these conditions, TOWA was not modulated by the time of day or circadian phase in various laboratory and nature-derived strains and did not correlate with baseline locomotor activity measured in standard activity monitors, indicating that it is distinct from spontaneous, clock-driven locomotion. Moreover, the finding that naïve TOWA did not decline in the first minutes during the analysis yet remained stable for at least 10 min argues against reactivity, which would be visible as a decrease in activity directly after introducing the fly to the arena. Instead, TOWA in the small arena appears to represent a form of stimulated activity ((Meehan and Wilson 1987), a term introduced to distinguish a stimulus-induced level of locomotor activity from spontaneous activity and reactivity during a reactivity period (Connolly 1967)). This stimulated activity is obviously modulated by aversive or appetitive stimuli, anxiolytic substance and manipulation of neuromodulatory signalling prior to the OFT, in line with the idea of basic emotion-like states in the fly. We assume that this stimulated activity may comprise elements of escape behaviour (one form of defensive behaviour). Interestingly, nature-derived strains showed higher TOWA than the highly inbred lab strains OrR and CS. This may not be surprising as OrR and CS flies have been selected for hundreds of generations for a sedentary lifestyle in fly bottles. In addition, compared to lab strains, nature-derived strains likely possess a higher sensory acuity and lowered arousal threshold. Collectively, our data suggest that TOWA in the OFT can be interpreted as a behavioural proxy for exogenously induced arousal. Moreover, our results showed that WAFO and TOWA are largely independent variables, which corresponds to the orthogonal conceptualisation of valence and arousal in dimensional theories of human emotions.

When we next plotted WAFO as an indicator of relative valence (i.e. fold change of – WAFO with respect to the pre-stimulus condition), and TOWA as a proxy of relative arousal (i.e. fold change of TOWA with respect to the pre-stimulus condition) in analogy to the circumplex model (Figure 7), most conditions and treatments lead to a location in the upper left Q4 quadrant that in humans is associated with conditions like “fearful”, “stressed” or “nervous”, or the lower right Q2 quadrant that in humans is typically associated with states like “relaxed”, “calm” or “contented” (Russell 1980; Posner et al. 2005; Mendl et al. 2010). Few conditions and treatments led to a location in the Q1-Q3 axis. Continued foot shocks as well as consumption of 10% ethanol led to a location in Q3 that in humans is associated with conditions like “depressed”, “bored” or “sad”. In comparison to a single electric foot shock that increased WAFO and TOWA, continued foot shock increased WAFO but slightly decreased TOWA. This phenotype is reminiscent of a depressive state in mammals, and resembles the learned helplessness behaviour of flies in the electric shock box in which uncontrollable electric shocks have a longer-lasting negative effect on locomotor activity (Batsching et al. 2016). Mating after rejection and sleep recovery showed the strongest decrease in WAFO with weakly increased TOWA and hence co-locate with a state that is labeled as “joy” in humans (Q1). We can currently only speculate why most conditions and treatments lead to a location along the Q2-Q4 axis, while only few lead to a location in the Q1-Q3 with a relatively small difference in TOWA compared to controls. However, we note that, in humans, the negative/inactive quadrant is also frequently underrepresented (Lang et al. 1993; Bradley and Lang 2000; Bradley et al. 2001), possibly because negative emotional states always induce some degree of arousal. Alternatively, the observed distribution may be caused by the nature of the OFT which is typically conceptualised as a behavioural assay for anxiety-like behaviour and thus promotes conditions along the Q2-Q4 axis which have been related to the avoidance of fitness-threatening punishers (Mendl et al. 2010). Moreover, the normalisation against control flies that have their major requirements satisfied (i.e., *ad libitum* access to food, mating and social partners present, optimal temperature and light regime) might negatively bias against emotional states along the Q1 (upper right) and Q3 (lower left) axis which have been related to acquiring fitness-enhancing rewards (Mendl et al. 2010).

**Figure 7.**
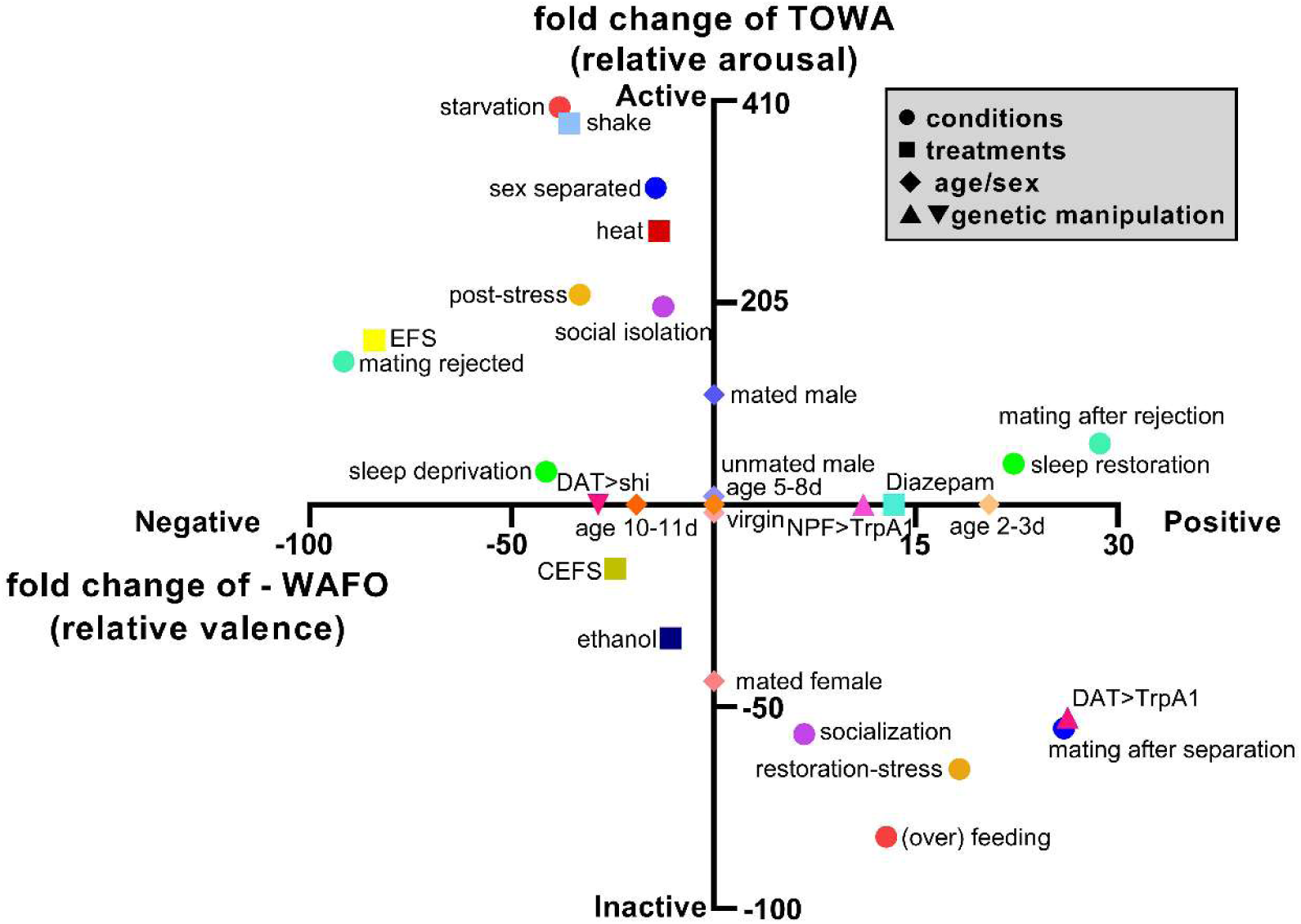
Summary of the results from all treatments plotted analogously to the circumplex model of valence and arousal. The median values from all treatments were normalized to the median control value: for post-stress, the pre-stress median served as the control, and for post-restoration, the post-stress median served as the control. These normalized values were then plotted as fold changes relative to the control state, with the fold change in -WAFO represented on the X-axis and the fold change in TOWA represented on the Y-axis. It is important to note that the ratio of the positive and negative axes on the XY plot is not consistent.

Taken together, plotting our WAFO-TOWA data in a circumplex-like system in Figure 7 produced a distribution that is remarkably similar to the distribution of emotional states in humans in the bipolar valence-arousal circumplex. In addition, the opposite effects of negative and positive stimuli on WAFO are compatible with the “principle of antithesis” (Darwin 1872) that states that opposite emotional states are bound to opposite behavioural expressions. Anderson and Adolph (2014) functionally defined four features common to different emotions across animals that serve as “building blocks” or “emotion primitives”: valence, scalability (intensity), persistence and generalisation. Above, we already argued that WAFO can serve as a proxy for valence. This is supported by a recent study in *Drosophila* that showed that chronic winning or losing in repetitive aggressive encounters induces internal states that are associated with opposite valence and affect other, non-aggressive behaviours such as gap-crossing or object exploration (Kim et al. 2018). The increasing changes in TOWA and WAFO upon extended duration of social isolation, sexual deprivation or mechanical shock suggest that the OFT behaviour is scalable with increasing stimulus strength. Further, the treatments (except for the electric shock) were in most cases given prior to the OFT. Thus, the observed behavioural effects in the OFT are in line with the assumption of a short-term persistent emotional-like state in the flies. In addition, different treatments led to a similar behavioural reaction. While this may largely be caused by the nature of the OFT set-up, it at least does not speak against the presence of a generalised response. Taken together, these findings lend behavioural support to the presence of basic emotion-like states in the fruit fly.

In mammals including humans, arousal is defined by autonomic nervous system activation with physiological consequences, e.g., sweating, changes in heartbeat, or pupil diameters that ideally are measured along with the behaviour. For example, in mice, measurement of thermal responses allowed to correlate physiological arousal with anxiety-related behavior (Lecorps et al. 2016), and the co-measurement of behaviour and heart rate dynamics allowed to refine and strengthen defensive states (Signoret-Genest et al. 2023). Importantly, changes in heart rate were also found to accompany defense behaviour in tethered *Drosophila* that were presented with a looming threat stimulus (Barrios et al. 2021). Heart rate decelerated during freezing, and accelerated during escape runs and a startle response upon looming (Barrios et al. 2021). A caveat of our study is that we do not provide physiological evidence for an aroused state since methods to measure heart rate in freely behaving flies are not available yet and were not developed in this study. To increase the validity of our conclusions, further studies are required that measure behaviour in combination with heart rate or other physiological parameters like metabolic rate or tracheal opening (Lehmann 2001).

### Manipulation of neurochemical signalling supports the presence of emotion primitives in Drosophila

The results obtained by genetic manipulation of neuromodulatory signalling pathways linked to human emotions provide further support for the presence of emotion primitives in the fly. Serotonin signalling has already been shown to influence WAFO in flies and rodents (Prut and Belzung 2003; Mohammad et al. 2016) and presents an ancient fear-related pathway (Mohammad et al. 2016). In contrast to Mohammad et al. (2016), both knockdown and overexpression of SerT lead to increased WAFO in this study. This non-linear response to serotonin signalling could be caused by level-dependent differential effects on the five serotonin receptor subtypes in the fly (Blenau 2026). For the levels of dopamine, the relationship with cognitive performance or working memory follows a U shape in primates (see e.g. (Desimone 1995; Cools and D’Esposito 2011). In *Drosophila*, both reduced and increased dopamine levels lead to increased male-to-male courtship behaviour (Liu et al. 2008, 2009). A similar situation may apply to the effect of serotonin in the OFT. In mammals, at least, serotonin levels either above or below an optimum level during development result in behavioural impairments in adults (see (Shah et al. 2018).

In both flies and mammals, dopamine and NPY/NPF signalling is associated with the reward system and hence positive valence (see (Dvořáček and Kodrík 2021)). Thermogenetic activation of these neuromodulator systems in the fly decreased WAFO, while inhibition of dopamine signalling increased WAFO.

Confirming a previous study (Mohammad et al. 2016), the anxiolytic GABA_A_ receptor blocker diazepam decreased WAFO, as expected from the anxiolytic effect of diazepam in humans. In rodents, diazepam mostly but not consistently leads to an anxiolytic response in the OFT behaviour which questions the usefulness of the OFT for testing anxiolytic drugs ((Prut and Belzung 2003; Rosso et al. 2022)).

Interestingly, inhibition of dopamine signalling, activation of NPF signalling and diazepam only affected WAFO, but not TOWA. This suggests the existence of at least partially distinct neuronal substrates for arousal and valence in the fruit fly, in line with the assumption that the two bipolar dimensions arousal and valence in the circumplex represent two independent neurophysiological systems (Russell 1980; Posner et al. 2005)

### Conclusions

Taken together, our results provide behavioural and neurochemical support to the idea that insects such as the fruit fly share stereotypical behavioural phenotypes and evolutionarily conserved and genetically hardwired neuronal circuits (“emotion primitives”) that underly basic emotions in humans (Gu et al. 2019). Moreover, the results provide additional support for the use of *Drosophila* as a genetically tractable model to study the neuronal substrate of “emotion primitives” and proposes the OFT in a small arena as an easy and automatable behavioural readout. Our study did not attempt to test whether flies have emotions or affective states linked to conscious feelings, nor are the results informative in this respect.

## Materials and Methods

### Fly strains

*Drosophila melanogaster* strains were cultured at 25°C and 60 % humidity on a 12 h light/ 12 h dark cycle on conventional cornmeal-yeast food. The following wild-type (WT) strains were used: CantonS (CS) Würzburg stock collection, CS^110^ that is CS crossed back to 110 generations to *per^01^* (Horn et al. 2019), Oregon R (OrR) Würzburg stock collection, *w*^1118^ (Bloomington Drosophila Stock Center (BDSC)), isogenic nature-derived WT strains Munich/Germany (coll. by John Parsch) and Akaa/Finland (coll. by Maaria Kankare) originated from the DrosEU project (Kapun et al. 2020), WT Hubland was collected close to the Biocenter in Würzburg (Deppisch et al. 2022). For the nature-derived WT strains, we reconstituted the population by outcrossing of isofemale lines. Transgenic/mutant flies used included *w*;tub-Gal80^ts^, yw;nSyb-Gal4*, w*;; *SerT^GD3824^* UAS-RNAi (Vienna Drosophila Resource Center (VDRC) #11346), *w*;* UAS-*SerT* (BDSC #24464), *w^1118^;GMR58E02-Gal4* (PAM*-Gal4,* BDSC #41347 (Liu et al. 2012)), *y^1^w*svr^PG33^/FM7^h^; hs-svr1A-2-3-t2* (BDSC #31411, 31413 (Sidyelyeva et al. 2010)), *y^1^w\**; *neuropeptide F* (*NPF*)-Gal4 (BDSC #25682), w*;UAS-*TrpA1*, and w*; UAS-*shibire^ts^*.

### Open field test (OFT)

Unless stated otherwise, 5-10 days old male CS flies were sedated on ice, then male flies were picked and individually placed into circular arenas with a diameter of 1 cm arranged in a 4×6 matrix drilled in a 2 mm thick dark-gray plastic plate. The arena plate was sandwiched between two glass plates to prevent flies from escaping, and placed on a white light plate (LED-Panel LP100-NW-Mw (Pollin)) with a light intensity of 270 lm within a climate chamber set to 20°C except for the experiments with *shi^ts^*flies which were conducted at 29°C. Before starting the recording system (sCMOS USB 3.0 Monochrome DMK 33UX287 camera, ImagingSource, Bremen, Germany), the arena plate was gently shaken to wake up the flies. Fly behaviour was recorded for 10 minutes at 5 frames per second (FPS). The trajectories of moving flies were automatically tracked using Ctrax (Branson et al. 2009). Dead or immobile flies were excluded from the analysis. The XY coordinates from Ctrax were then further analysed using a custom-made Matlab script (developed by YLW and Stefanie Pütz) for total-walking distance (TOWA). Wall-following behaviour (WAFO) was calculated as the distance to the center of the arena as calculated from the XY coordinates.

### Conditions and treatments prior to the OFT

#### Social isolation

After a control OFT, male flies were separated and kept individually in a single vial on food for 4 days without any social interaction, with OFT after day 2 and day 4 of isolation. Afterwards, the male flies were pooled in one vial, and the same number of females were added, followed by a last OFT on the next day.

#### Starvation

After a control OFT, male flies were maintained for 2 days on agarose to provide water but no food source, with OFT after day 1 and 2 of starvation. Afterwards, flies were pooled in one vial containing standard food, followed by a last OFT the next day.

#### Sleep deprivation

After a control OFT, male flies were placed at ZT3 on a vibrating shaker to prevent sleep. At ZT7 (after siesta) and ZT13 (beginning of the night), flies were re-analysed in the OFT. Afterwards, the need for sleep was restored by placing the flies overnight on a stable surface before the measurement in last OFT.

#### Sex deprivation

We used two different sex-deprivation schemes to test the effect of a lack of courtship in male flies:

1. Males were kept in a mixed-sex group for seven days and then tested in the OFT to obtain the pre-stress control value. Then, the males were separated from females and maintained in a male-only group for 2 days, with OFT after day 2 and day 4 of isolation. Afterwards, males were pooled in one vial with the same number of virgin females, followed by a last OFT on the next day.
2. Males were collected directly after eclosion and kept as male-only group for two days, to obtain the pre-stress control value. Then, the males were paired for one hour with a similar number of females that had been mated on the previous day, and courtship behaviour was recorded. Males that were rejected by the mated female during that hour were then tested in the open field on the next day. Pairing and rejection was repeated on the next day, followed by a third OFT on the next day. Afterwards, males were pooled in one vial with the same number of virgin females, followed by a last OFT on the next day.

#### Multiple stressors

After a control OFT, male flies were subjected to a strong stress situation for 24 h that combined social isolation, starvation and sleep deprivation as described above, followed by an OFT. Then, the males were pooled in one vial containing normal food without agitation, and an equal number of virgin females was added, followed by a last OFT on the next day.

#### Mechanical shock

After a control OFT, the arena with the flies was five times sharply tapped on the table edge, directly followed by an OFT. This was followed by a second round of tapping and recording as described.

#### Heat shock

After a control OFT at the standard temperature of 20°C, the arena was placed on a heat plate and heated to 37°C directly followed by a second OFT.

#### Electric foot shock (EFS) and consecutive EFS (CEFS)

*EFS*: after a control OFT, a custom-made plate with an electric grid was inserted between the glass floor and the arena wall. Then, flies received an EFS a 90V EFS for 1.2 seconds which was repeated 10x with an interval of 1 minute. The strength of the EFS was strong but not deleterious according to a pre-evaluated response curve. Then, the electric grid was removed and a second OFT was conducted.

*CEFS*: first, male flies were treated with an EFS as described above. Next, the males were transferred back to a vial containing a similar number of females on standard food. On the next day, flies were measured in the OFT, and then received a consecutive EFS, followed by another OFT.

### Pharmaceutical treatments

#### Ethanol

For assessing the effects of ethanol, a Capillary Feeder (CAFE) system (Ja et al. 2007) with 5 μl capillaries was used to provide liquid food containing 10% ethanol, or pure water as control. The system consisted of a plate with 24 wells tiled with moist filter paper and closed by a lid was used to house flies individually. Capillaries were inserted into the well through small holes drilled into the lid and an underlying layer of rubber. Test capillaries in wells without flies served to correct for evaporation. On the first training day, the experimental and the control group were fed with freshly prepared food made of 5.4% sucrose in water. The well plate was placed in an incubator at 25 °C and 60 % humidity in a 12 h light/ 12 h dark cycle for approximately 24 hours. On the second day, flies were maintained in the wells, but food-containing capillaries were swapped against water-only filled capillaries for approximately 24 hours to promote hunger. On the third day, the control group was given the same freshly prepared 5 μl 5.4% sucrose water solution as on the training day in the CAFE system. The experimental group received 5 μl 10% ethanol in 5.4% sucrose water solution. Both groups were placed in the incubator for the next 24 hours under the same conditions as above. Finally, flies that consumed at least 1 µl were analysed in the OFT to assess the ethanol effect on open field behaviour.

#### Diazepam

Male flies were starved overnight in 2% agarose-only vials for 17 hours at 25°C and 60 % humidity on a 12 h light/ 12 h dark cycle. Afterwards, control flies were transferred to a 7 ml-volume food vial with 4% sucrose and 2% agarose. The experimental group was maintained in the food vial with 5 mM diazepam in 4% sucrose and 2% agarose. A red food dye was added to both foods, and only flies with colored abdomens were collected for conducting a subsequent OFT.

#### Thermogenetics

For temperature-controlled expression of *dSerT and dSert-RNAi*, the TARGET system (McGuire et al. 2004) was used, based on the temperature-sensitive driver line *Tub-Gal80^ts^, nSyb-Gal4* which was crossed with either *UAS*-*dSerT* or *UAS-dSert-RNAi* effector lines. For temperature-controlled inhibition or activation of dopaminergic and NPF neurons, the *GMR58E02-* and the *NPF-Gal4* driver line were crossed to either *UAS-TrpA1* and *UAS-shibire^ts^* effector line. TRPA1 is a temperature-dependent cation channel that opens at temperatures around 29°C and induces neuronal activation (Rosenzweig et al. 2005). *shibire^ts^* is a temperature-sensitive dynamin that loses its function for synaptic vesicle recycling at temperatures around 29°C or higher (Kitamoto 2001).

The newly eclosed progeny one of the respective Gal4xUAS crosses were raised at 18°C for 2–3 days prior to the behavioural assays. Then half of the animals were transferred to 29°C overnight (crossing with *UAS-TrpA1* and *UAS-shibire^ts^*) or three consecutive days (crossing with *Tub-Gal80^ts^, nSyb-Gal4*) before the OFT, while the other half were kept at 18°C. On the test day, all flies were held at 25°C for a 3 h recovery period prior to the assay. Like the animals in experimental group, animals as genetic control raised at identical conditions were generated by crossing the driver line with *w^1118^*and the effector line with *w^1118^*.

To generate peptide-depleted flies, *y^1^ w* P{w^+mWhs=GawB^}svr^PG33^/FM7^h^; hs-svr^1A-2-3-t2^* flies (Sidyelyeva et al. 2010) were heat-shocked every day for 30 minutes to rescue the loss of carboxypeptidase D (SILVER), a key enzyme required for neuropeptide biosynthesis (Pauls et al. 2019). Three to five days after eclosion, the heat shock was discontinued for 1-3 days prior to the OFT to stop bioactive peptide production.

#### Locomotor activity assay

The locomotor activity of 2-3 days old male post eclosion, was recorded with the *Drosophila* Activity Monitor (DAM) system (Trikinetics, Princeton, MA, USA). 28 flies were placed in separate glass tubes with a diameter of 0.5 cm. The tube was filled on one end with food consisting of 4% sucrose and 2% agar in water. This end was sealed with a rubber plug, while the open end was blocked by a sponge plug. The activity was recorded for eight days under LD12:12 cycles at 20°C. Data from day 2 to 8 recording were utilized to calculate the mean locomotor activity at ZT0, ZT6, ZT12, and ZT18. To enhance comparability with the OFT, the mean activity was binned in 10-minute intervals during ZT0-1, ZT6-7, ZT12-13, and ZT18-19. This calculation was performed over a one-week period for 28 individual flies, and the results were plotted as activity counts per number of times.

## Data analysis and statistics

Custom-made Matlab scripts were used to analyze the open field behaviour. The arena was seen as a coordinate system with an x- and y-axis, and the coordinates of the center of the arena were defined as:

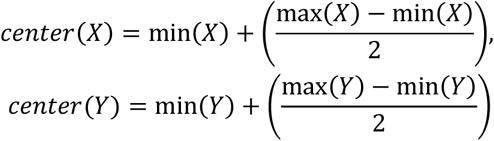

With the instantaneous coordinates of the fly (X_i_, Y_i_) and the center coordinates, wall-following (WAFO) of the fly was calculated in pixels by applying the Pythagorean theorem:

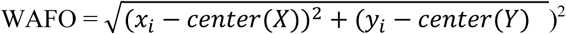

With the instantaneous coordinates of the fly (X_i_, Y_i_) and the coordinates on the next frame (instant location i+1), total walking activity (TOWA) of the fly during the 10 min recording was calculated in pixels by applying the summation:

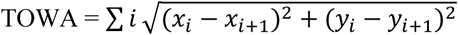

For the analysis, the median value of the WAFO in each frame during the 10-minutes at 5 fps recording was calculated. According to the ratio between the diameter of the arena in mm and in pixel, WAFO and TOWA were converted from pixel to mm. Flies that did not move all along the wall once were excluded from the analysis as our method does not allow to correctly calculate the center of the arena in that case.

The statistical evaluation and diagrams plotting were carried out by GraphPad PRISM 8.0 (Dotmatics), and by R 4.3.1 (www.r-project.org) using a custom-made script kindly provided by Morgan Oberweiser (University of Greifswald, Germany). For two group comparisons, depending on the data distribution, either an unpaired t-test or a Mann-Whitney U test was used. When more than two groups were compared to each other, either a one-way ANOVA (parametric data) or a Kruskal-Wallis test (non-parametric data) was applied.

## Acknowledgments

We thank Angelika Erhardt-Lehmann (University Hospital Würzburg) for help with the pharmacological experiments, Matthias Gamer (University of Würzburg) and Angelika Erhardt-Lehmann for valuable comments on the manuscript draft, the late Konrad Öchsner (University of Würzburg) for help in designing and setting up of the OFT setup, the Biocenter workshop for building the OFT setup, Maaria Kankare (University of Yyväskylä) and John Parsch (LMU Munich) for the kind gift of flies, Morgan Oberweiser (University of Greifswald) for her generous sharing of R codes for data visualisation, and Jan Ache (University of Würzburg) for helpful discussion regarding data presentation. The work was made possible by funding through the Deutsche Forschungsgemeinschaft (DFG, WE 2652/9-1) to CW.

We dedicate this paper to the memory of Konrad Öchsner.

## Author contributions

Conceptualization: YLW, CW; Methodology: YLW, CW; Formal analysis: YLW, ED, HW, CW; Investigation: YLW, ED, HW, MS; Writing and Visualisation: YLW, HW, CW; Writing review: ED, MS; Funding acquisition: CW.

## Supplementary table

**Suppl. Table 1:** Statistics for Figure 5A-B. The data for the different ZT times were analysed by a Kruskal-Wallis test. If significant differences were found, Dunn’s multiple comparison test was used for pairwise comparison between ZT times.

| Oregon R |  |  |  | CS |  |  | CS<sup>110</sup> |  |  | Munich |  |  | Hubland |  |  | Finland |  |  |
| --- | --- | --- | --- | --- | --- | --- | --- | --- | --- | --- | --- | --- | --- | --- | --- | --- | --- | --- |
| WAFO (mm) | mean | sd | n | mean | sd | n | mean | sd | n | mean | sd | n | mean | sd | n | mean | sd | n |
| ZT0 | 2.61 | 0.71 | 23 | 2.60 | 0.54 | 22 | 3.25 | 0.50 | 24 | 3.30 | 0.38 | 21 | 3.37 | 0.48 | 22 | 3.62 | 0.49 | 24 |
| ZT6 | 2.33 | 0.53 | 22 | 2.64 | 0.56 | 22 | 3.26 | 0.34 | 21 | 3.24 | 0.31 | 22 | 3.49 | 0.31 | 24 | 3.51 | 0.21 | 23 |
| ZT12 | 2.68 | 0.79 | 23 | 2.48 | 0.55 | 22 | 3.57 | 0.53 | 24 | 3.21 | 0.26 | 23 | 3.61 | 0.26 | 22 | 3.80 | 0.41 | 24 |
| ZT18 | 2.49 | 0.86 | 22 | 2.70 | 0.68 | 24 | 3.60 | 0.62 | 24 | 3.45 | 0.38 | 23 | 3.39 | 0.45 | 22 | 3.60 | 0.38 | 23 |
| significance | ns |  |  | ns |  |  | ZT6 vs.18: p=0.045 |  |  | ns |  |  | ns |  |  | ns |  |  |
| TOWA (m) | mean | sd | n | mean | sd | n | mean | sd | n | mean | sd | n | mean | sd | n | mean | sd | n |
| ZT0 | 0.17 | 0.28 | 23 | 0.39 | 0.33 | 22 | 1.03 | 0.53 | 24 | 0.91 | 0.57 | 21 | 1.43 | 0.73 | 22 | 1.10 | 0.63 | 24 |
| ZT6 | 0.13 | 0.06 | 22 | 0.52 | 0.35 | 22 | 0.97 | 0.41 | 21 | 1.20 | 0.54 | 22 | 1.71 | 0.59 | 24 | 2.07 | 0.61 | 23 |
| ZT12 | 0.16 | 0.16 | 23 | 0.39 | 0.37 | 22 | 0.76 | 0.34 | 24 | 1.19 | 0.70 | 23 | 1.79 | 0.54 | 22 | 1.24 | 0.61 | 24 |
| ZT18 | 0.17 | 0.31 | 22 | 0.50 | 0.39 | 24 | 0.66 | 0.47 | 24 | 1.05 | 0.53 | 23 | 1.34 | 0.62 | 22 | 1.55 | 0.70 | 23 |
| significance | ns |  |  | ns |  |  | ZT0vs.ZT18: p=0.024 |  |  | ns |  |  | ns |  |  | ZT0vsZT6: p<0.0001<br>ZT0vs.ZT6: p=0.002<br>ZT0vs.ZT18: p=0.037 |  |  |

## Supplementary figures

**Figure 1- figure supplement 1.**
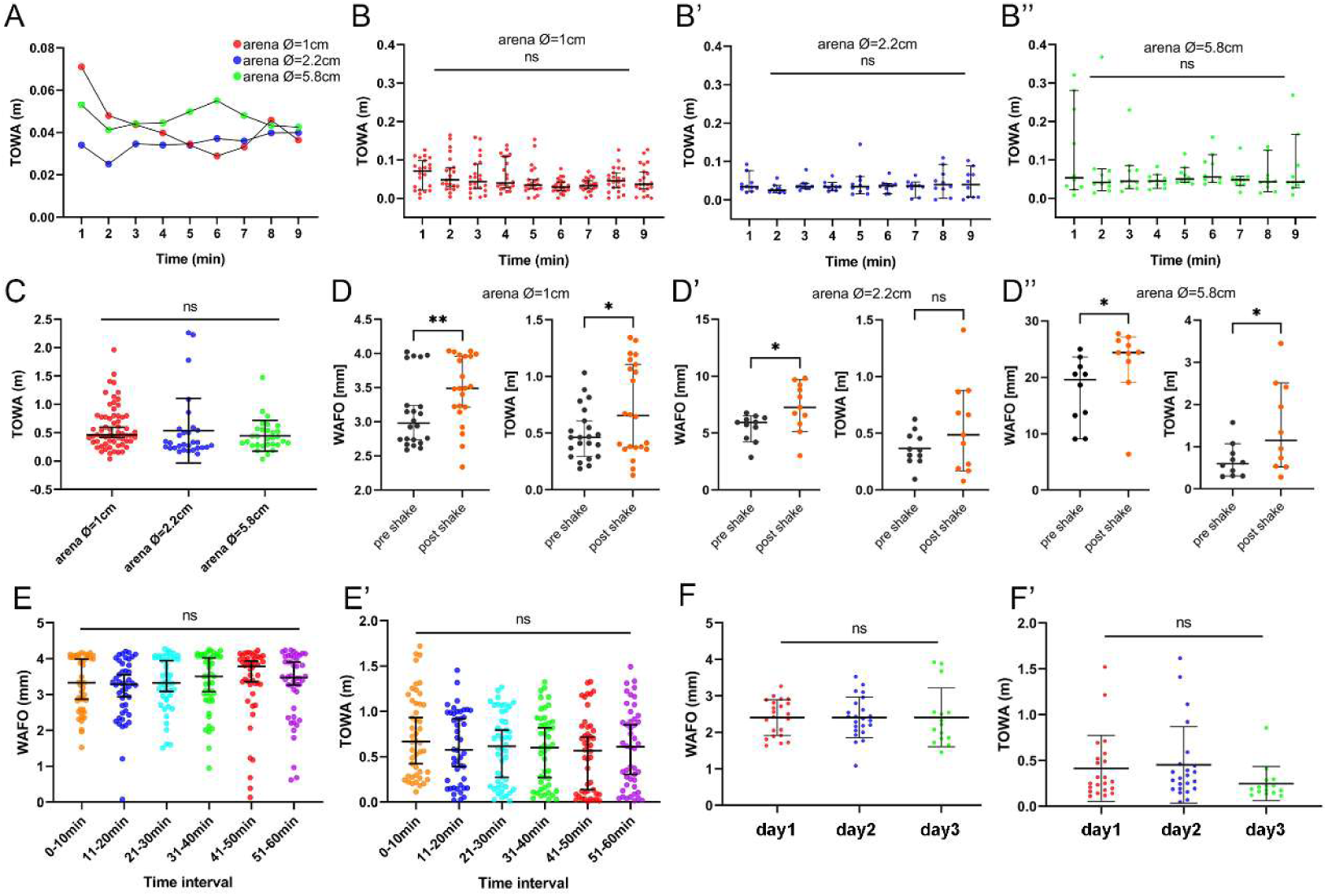
Open field behaviour of CantonS flies in circular arenas with a diameter of 1.0, 2.2 or 5.8 cm. **A)** Mean TOWA per minute in naïve flies was quite constant during the first 10 min of recording. **B-B’’)** Statistics for the TOWA data in A) for the arena diameter of **B)** 1 cm, **B’)** 2.2 cm and **B’’)** 5.8 cm. Although there was a trend towards higher activity in the first minute in the 1 cm arena, the difference is not significant.**C)** Accumulated TOWA during the 10 min in B-B **’**) did not reveal statistically significant differences between the arena sizes. **D-D’’)** WAFO and TOWA responses to mechanical stress (shake by tapping) in the arenas with different diameters. In all arena sizes, flies showed significantly increased WAFO after tapping. TOWA was similarly increased, yet the increase is not statisticially significant for the 2.2 cm arena. **E-E’)** WAFO and TOWA in 10 min bins did not vary significantly during a 60 min recording of naïve untreated flies. **F-F’)** The same set of naïve flies were tested once per day in the OFT on three consecutive days. WAFO and TOWA was similar on all three days, suggesting that flies do not habituate to the situation.

**Figure 1- figure supplement 2.**
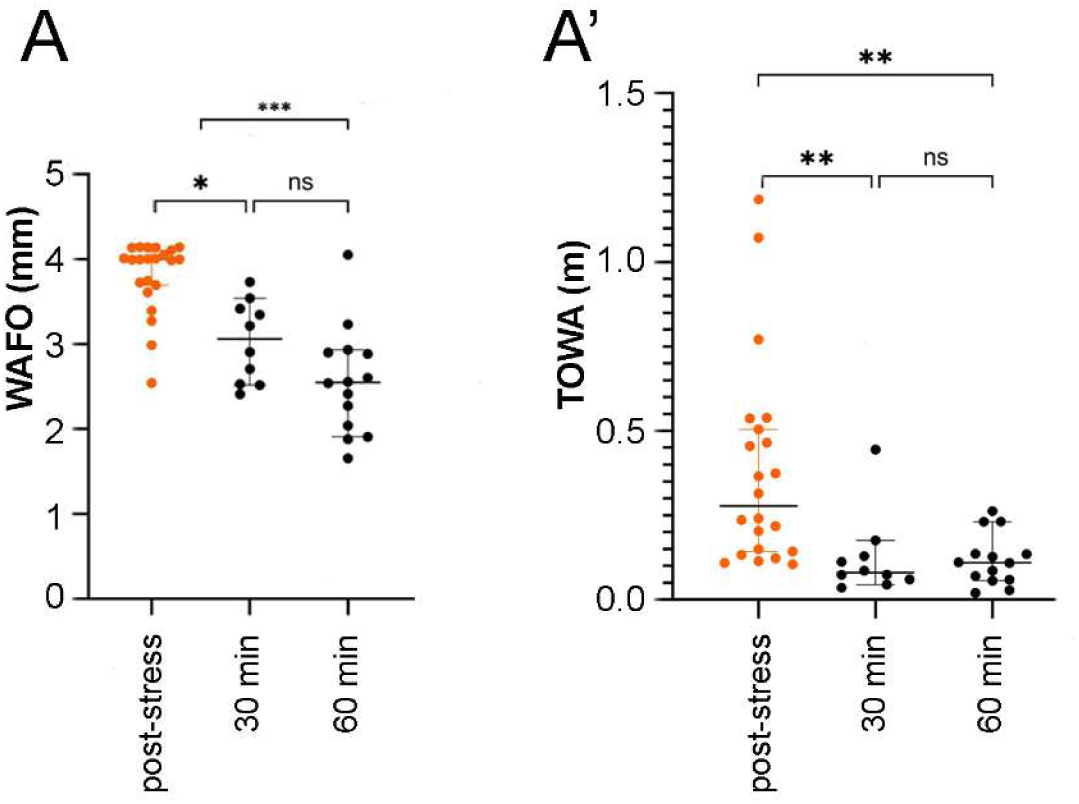
Open field behaviour of CantonS flies after mechanical stress (shake) in the 1cm arena. **A)** Mean WAFO directly after mechanical stimulus (tapping, orange dots post-stress), 30 min or 60 min later. It is visible that the stress-induced WAFO is only temporary and has significantly declined to the level of naïve flies after 30 and 60 min. **A’)** Mean TOWA directly after mechanical shake (orange dots, post-stress), 30 min or 60 min later. It is visible that the stress-induced increase in TOWA is only temporary and has significantly declined to the level of naïve flies after 30 min which remains constant until 60 min post-stress.

**Figure 4-figure supplement 1.**
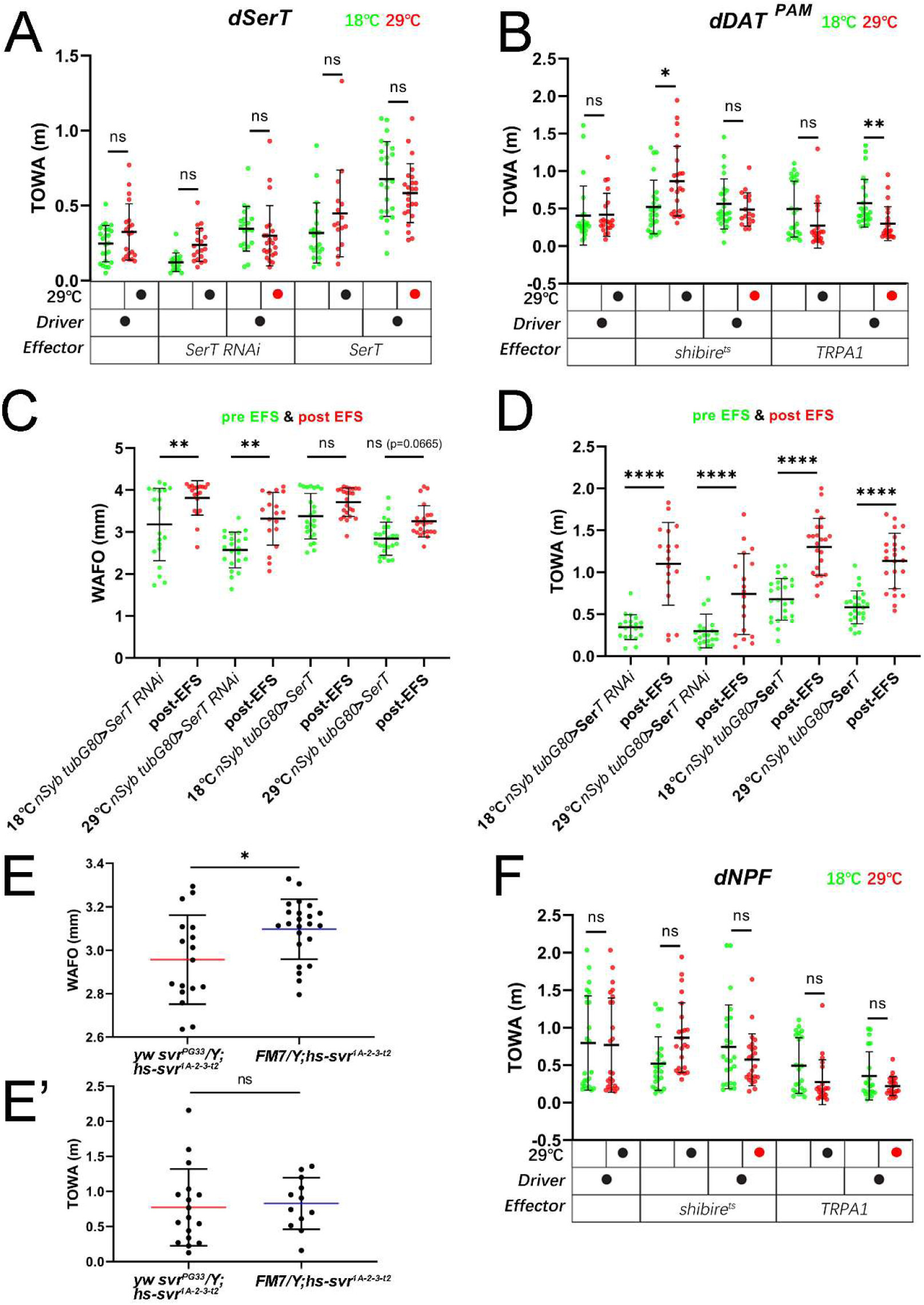
Effects of genetic manipulation of biogenic amine and neuropeptide signalling on open field behaviour. **A)** Flies with temperature-induced (29 °C, red) conditionally reduced serotonin transporter expression (nSyb tubGal80^ts^ x SerT RNAi) or UAS-mediated SerT overexpression (nSyb tubGal80^ts^ x SerT) showed no change in TOWA compared to flies at the non-inducing condition (18 °C, green). **B)** Thermogenetic suppression of dopaminergic reward neurons of the PAM cluster by a temperature-sensitive shibire allele did not alter TOWA compared to the controls at 18°C. Thermogenetic activation of the same PAM neurons by TRPA1 decreased TOWA compared to the temperature controls. **C)** Flies with reduced serotonin transporter expression (nSyb tubGal80^ts^ x SerT RNAi) cultured at 18°C or 29°C before receiving EFS (green) and after receiving EFS (red) showed a significant difference in WAFO. Flies with UAS-mediated SerT overexpression (nSyb tubGal80^ts^ x SerT) showed a trend of increased WAFO after receiving EFS. **D)** Flies with reduced serotonin transporter expression (nSyb tubGal80^ts^ x SerT RNAi) and flies overexpressing serotonin transporter (nSyb tubGal80^ts^ x SerT) cultured at 18°C and 29°C all showed a significant increased TOWA after receiving EFS (red) compared to the state before receiving EFS (green). **E)** WAFO and **E’)** TOWA in flies with conditionally blocked neuropeptide production (svr^PG33^ mutants) compared to controls (FM7). Blocked neuropeptide production for 1-3 days resulted in decreased WAFO (E), while TOWA remained unaffected (E’). **F)**Thermogenetic suppression and activation of NPF-positive neurons had no effect on TOWA compared to the respective temperature controls.

